# AMPing up the search: An *in silico* approach to identifying Antimicrobial Peptides (AMPs) with potential anti-biofilm activity

**DOI:** 10.1101/2021.08.16.456477

**Authors:** Shreeya Mhade, Stutee Panse, Gandhar Tendulkar, Rohit Awate, Snehal Kadam, Karishma S Kaushik

**Affiliations:** Guru Nanak Khalsa College of Arts, Science and Commerce (Autonomous), Mumbai, India; The Pennsylvania State University, University Park, Pennsylvania, USA; Sir Sitaram and Lady Shantabai Patkar College of Arts and Science and V P Varde College of Commerce and Economics (Autonomous), Mumbai, India; Northeastern University, Boston, USA; Hull York Medical School, University of Hull, United Kingdom; Department of Biotechnology, Savitribai Phule Pune University, Pune, India

**Keywords:** Biofilms, antimicrobial peptides, sortase C, *C. striatum*, protein-peptide molecular docking

## Abstract

Antibiotic resistance is a public health threat, and the rise of multidrug-resistant bacteria, including those that form protective biofilms, further compounds this challenge. Antimicrobial peptides (AMPs) have been recognized for their anti-infective properties, including their ability to target processes important for biofilm formation. However, given the vast array of natural and synthetic AMPs, determining potential candidates for anti-biofilm testing is a significant challenge. In this study, we present an *in silico* approach, based on open-source tools, to identify AMPs with potential anti-biofilm activity. This approach is developed using the sortase-pilin machinery, important for adhesion and biofilm formation, of the multidrug-resistant, biofilm-forming pathogen *C. striatum* as the target. Using homology modeling, we modeled the structure of the *C. striatum* sortase C protein, resembling the semi-open lid conformation adopted during pilus biogenesis. Next, we developed a structural library of 5544 natural and synthetic AMPs from sequences in the DRAMP database. From this library, AMPs with known anti-Gram positive activity were filtered, and 100 select AMPs were evaluated for their ability to interact with the sortase C protein using *in-silico* molecular docking. Based on interacting residues and docking scores, we built a preference scale to categorize candidate AMPs in order of priority for future *in vitro* and *in vivo* biofilm studies. The considerations and challenges of our approach, and the resources developed, which includes a search-enabled repository of predicted AMP structures and protein-peptide interaction models relevant to biofilm studies (B-AMP), can be leveraged for similar investigations across other biofilm targets and biofilm-forming pathogens.

## Introduction

The spread of multidrug-resistant bacteria, and the emergence of pathogens inherently resistant to a range of antibiotics, is a serious challenge. This problem is further exacerbated by the ability of bacteria to form protective communities or biofilms. In biofilms, bacteria are notoriously tolerant to antibiotics and often require long-term treatments, which further fuels the development of antibiotic resistance. Given this, there is an urgent need to search for novel anti-biofilm approaches to complement and potentially serve as alternatives to standard antibiotics. Antimicrobial peptides (AMPs) are a diverse class of peptides with a range of structural and functional properties [1]. In addition to their well-known effects on disrupting bacterial membrane integrity, several AMPs target specific bacterial components, and can thereby inhibit processes such as biofilm formation [2]. Bacterial cell-surface pili play an important role in the various stages of biofilm formation, including initial attachment and adhesion [3]. In Gram-positive bacteria [4], pilus biogenesis is mediated by a group of sortase enzymes, and inhibition of sortase activity results in defective biofilm formation [5–7]. AMPs have been shown to demonstrate activity as sortase inhibitors, with the potential to act in adjunct with standard antibiotics [8–10]. While this is promising, given the vast array of AMPs, identifying potential candidates for anti-biofilm testing remains a significant challenge.

In this study, we present an *in silico* approach to identify AMPs with a potential anti-biofilm activity using the sortase-pilin machinery of the multidrug-resistant, biofilm-forming pathogen *Corynebacterium striatum* as a target. Using homology modeling, we modeled the semi-open lid conformation of the *C. striatum* sortase C protein adopted during pilus biogenesis. With a combination of predictive peptide modeling tools, we developed structural models of 5544 natural and synthetic AMPs from the DRAMP database. From this library, AMPs with known anti-Gram positive activity were filtered, and 100 select AMPs were evaluated for their ability to interact with the sortase C protein using *in-silico* molecular docking. Docking results were used to build a preference scale to categorize AMPs in order of priority for future testing against *C. striatum* biofilms. In doing so, this *in silico* approach enables the identification and categorization of candidate AMPs with potential anti-biofilm activity, and provides a starting point for *in vitro* and *in vivo* biofilm testing. Finally, while our study focuses on *C. striatum*, we argue that the approach we present, considerations described and resources developed, which includes a search-enabled repository of predicted AMP structures and protein-peptide interaction models for biofilm studies (Biofilm-AMP or B-AMP), can be leveraged for similar investigations across other biofilm targets and biofilm-forming pathogens.

## Methods

### Homology modeling of the *C. striatum* sortase C protein

The three-dimensional structure of the sortase C protein of *Corynebacterium striatum* is not available in the RCSB Protein Data Bank (PDB) [11]. To construct the protein structure of the *C. striatum* sortase C enzyme, the amino acid sequence of the sortase C protein was retrieved from the GenPept Database (ID: WP_034656562) [12]. The physicochemical properties of the protein such as molecular weight, isoelectric point (pI), amino acid composition, estimated half-life, and instability index were determined using the ProtParam tool [13]. The secondary structure of the sortase C protein was predicted by the PSIPRED server [14]. Both physicochemical and secondary structure properties (helix, strand, or coil) of the protein were determined using the amino acid sequence as an input. The three-dimensional (3D) protein structure of the class C sortase enzyme was constructed using I-TASSER [15], by submitting the FASTA sequence retrieved from GenPept as an input (without any user-specific restraints). To mimic the semi-open lid conformation of the *C. striatum* sortase C protein, we generated a model that included a double mutation D93A/W95A (replacing Aspartate (93) and Tryptophan (95) with Alanine) in the protein. The input FASTA file was modified manually by replacing A in place of D and W.

### Validation of the predicted 3D model of the *C. striatum* sortase C protein

The reliability of the sortase C protein model, including the modified protein model with semi-open lid conformation (generated with I-TASSER), was verified using Structure Analysis and Verification Server version 6.0 (SAVES) [16–18]. The SAVES 6.0 server integrates analysis from multiple, widely-used validation algorithms (including VERIFY 3D, ERRAT, and PROCHECK). The PDB file generated from I-TASSER was submitted as an input to SAVES 6.0. The quality and reliability of the generated model were checked by using the default quality parameters of the ERRAT and VERIFY3D software. In addition, the PROCHECK software was used to validate the generated model of the proteins by utilizing the Ramachandran plot.

### Predictive modeling of AMP structures from the DRAMP database

From DRAMP V3.0 [19], we downloaded a dataset of 5562 general AMP sequences, with known antibacterial and antifungal activities. These AMPs were listed as Pep2 to Pep5563, where Pep1 is the pilin subunit (GenPept ID: WP_170219081.1) [20] **(Suppl File 1).** FASTA files were generated using an *in-house* developed Python script. The Python script reads the .csv file from the DRAMP database and generates FASTA files which were used for predictive modeling.

#### Peptide modeling with PEP-FOLD3

For modeling AMPs with less than 50 amino acid residues, we used PEP-FOLD3 [21], which predicts 3D peptide structures from sequence information using a *de novo* approach. FASTA file for the structure to be generated was uploaded in the query form, following which a peptide structure for the given sequence was generated and saved in .pdb form.

#### Peptide modeling with Robetta

For modeling AMPs with more than 50 amino acids, we used the deep-learning-based method TrRosetta [22] available under Robetta [23]. FASTA files of the peptide were uploaded in the query submission interface and the resulting model was downloaded with a .pdb extension.

#### Peptide modeling using ab-initio I-TASSER modeling

For peptides with unusual or unknown (X) amino acids, we used the *ab-initio* modeling feature of I-TASSER [15,24] for generating 3D conformations. FASTA files of the peptide sequence were uploaded into the query submission form using the default parameters, and the resulting models were downloaded with a .pdb extension.

### Protein-peptide *in silico* molecular docking of predicted AMP structures to the mutated *C. striatum* sortase C protein

For protein-peptide *in silico* molecular docking, we filtered out AMPs with annotated anti-Gram positive activity (based on information available in the DRAMP database) using an *in-house* developed Python script (https://github.com/KarishmaKaushikLab/B-AMP). The Python script employs a conditional statement to automatically scan the .csv file for the term “Anti-Gram+” from the activity column in the sheet **(Suppl File 2)**. From this filtered subset, 88 AMPs ranging in length from 2 to 8 amino acid residues, and 12 representative peptides with lengths ranging from 9 to 20 residues were selected for *in-silico* molecular docking. Before molecular docking to the mutated sortase C protein, the 3D AMP structures were subject to Energy Minimization in Swiss-PDBViewer SPDBV v4.1.0 [25] using the partial implementation of the GROMOS96 43B1 force field [26,27]. As a standard, the LPMTG motif of the pilin subunit was docked to the mutated sortase C protein. AutoDockTool1.5.6 [28] was used to generate the necessary input PDBQT files for docking, and AutoDock Vina [29] was used to performing protein-peptide docking on an Intel^®^ Core™ i7-975OH CPU @ 2.60GHz system using the command: vina --receptor M_Ala_Sortase.pdbqt --ligand Pep#.pdbqt --center_x 57.854 --center_y 47.915 --center_z 46.936 --size_x 28 --size_y 28 --size_z 28 --out vPep#.pdbqt --log vPep#.log. The run time for each docking varied depending on the length of the peptide and ranged from 2 minutes (for peptides 2-4 residues long) to 40 minutes (for peptides 20 residues long) with the system used.

### Building a structural repository of AMPs and protein-peptide interactions for biofilm studies (B-AMP)

To enable the availability of resources and data generated in this study, we developed a user-friendly, and search-enabled repository of AMP structures and AMP interactions with biofilm targets (B-AMP) (https://b-amp.karishmakaushiklab.com/). The repository was built using HTML/CSS/JavaScript delivered over GitHub pages, and the scripts and other resources employed have been made available (https://github.com/KarishmaKaushikLab/B-AMP).

## Results and Discussion

### Sortase C as an anti-biofilm target

Sortase enzymes function to assemble pili or anchor them to the bacterial cell wall, and have been identified as promising antimicrobial targets, particularly in Gram-positive bacteria [30–33]. Across various species, inactivation of the genes encoding sortase enzymes, including sortase C, has been shown to result in a significant reduction in biofilm formation [6,34–36]. Given this, sortases have been recognized as a promising target for anti-biofilm approaches [37]. In *C. striatum*, cell-surface pili facilitate adhesion to host cells and abiotic surfaces, and mediate the formation of biofilms [3]. The sortase-pilin machinery includes the *spaDEF* operon that encodes a set of pilus proteins and their respective sortases [38,39]. In *Corynebacteria* spp, the sortase C enzyme functions as a pilus-specific sortase, recognizing and cleaving the sorting signal (LPMTG) in the pilin subunits, which is followed by their surface display [30,33]. Based on its relevance to *C. striatum* pilus biogenesis, the sortase C protein was selected as the anti-biofilm target.

### Sequence-based prediction of physicochemical properties and secondary structure of the *C. striatum* sortase C protein

The amino acid sequence of the *C. striatum* sortase C protein was retrieved from the GenPept Database (ID: WP_034656562) [12]. Using the sequence, the physicochemical properties of the protein were predicted using ProtParam analysis. As shown in **Table 1,** the analysis revealed a molecular weight of 35134.72 Da, and a theoretical isoelectric point of 5.38 and 5.46 respectively, indicating a negatively-charged protein. In particular, the sortase C protein contained 46 negatively charged residues (Asp+Glu) and 33 positively-charged residues (Arg+Lys). The computed instability index of 33.54 indicates that the protein is stable, as the obtained value is less than the cut-off of 40. The N-terminal sequence containing M (Met) indicates that the half-life of the protein is 30 hours (mammalian reticulocytes, *in vitro*), >20 hours (yeast, *in vivo*), >10 hours (*Escherichia coli, in vivo*). Using the PSIPRED server [14], the secondary structure of the sortase C protein was observed to contain a combination of alpha-helices (41%), coils (44%), as well as beta sheets (15%) **(Suppl Figure 1).**

**Table 1.**
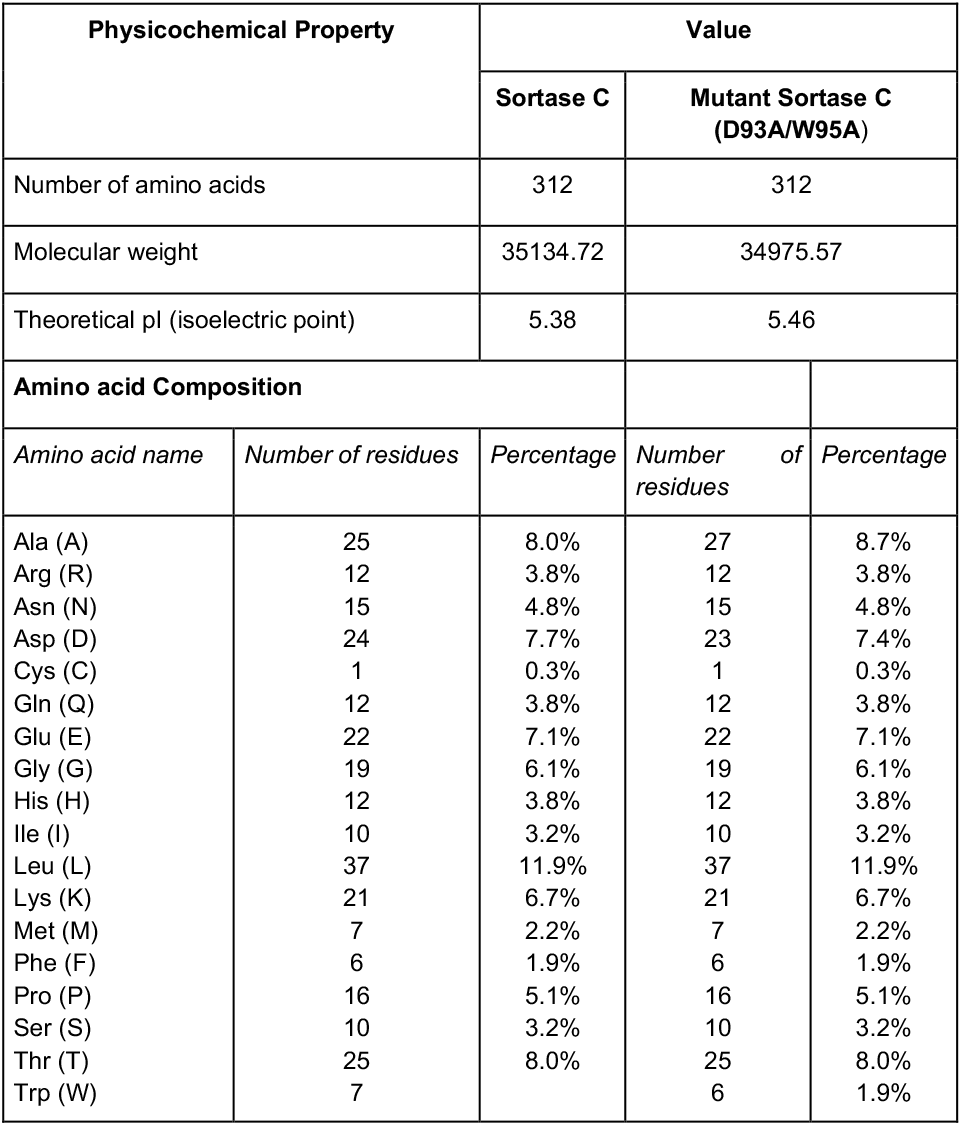
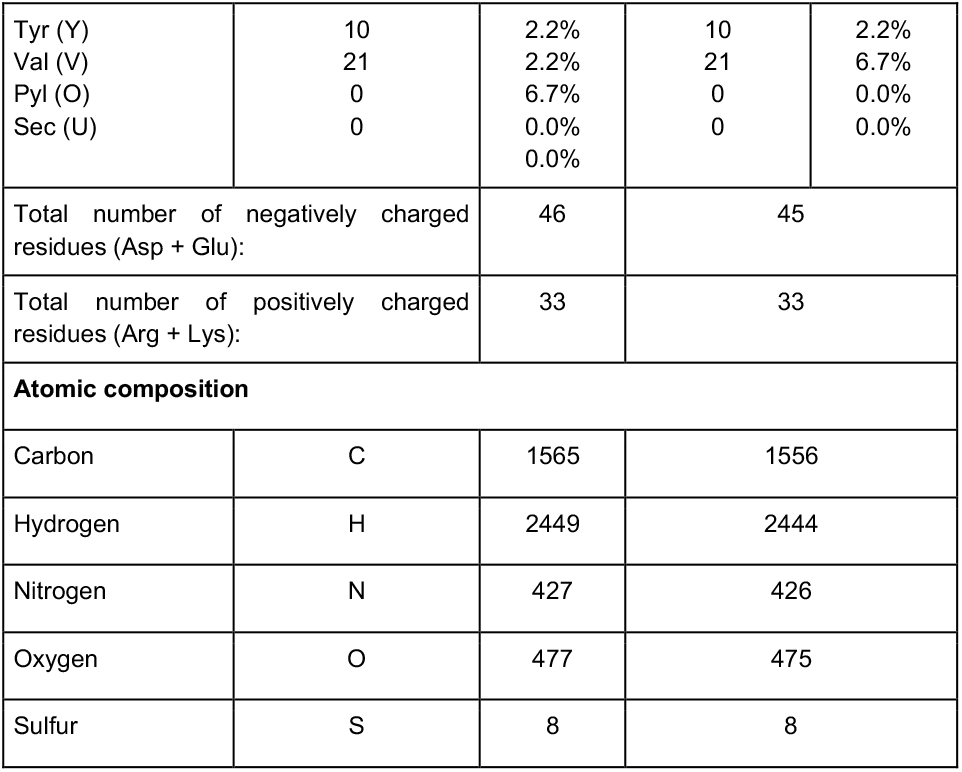
Physicochemical properties of the *C. striatum* sortase C protein (wild-type and double mutant D93A/W95A) using ProtParam analysis.

### Homology model of the *C. striatum* sortase C protein

The three-dimensional structure of the sortase C protein of *C. striatum* is not available in the RCSB Protein Data Bank (PDB) [11]. To model the structure of the enzyme, I-TASSER selected the following crystal structures as templates: class C sortases of *Streptococcus pneumoniae* (PDB ID: 2W1J; 2W1K; 3G66; 3O0P), *Streptococcus agalactiae* (PDB ID: 3RBI; 4D7W; 4G1H), *Actinomyces oris* (PDB ID: 2XWG), *Clostridium perfringens* (PDB ID: 6IXZ) and Class A sortase from *Corynebacterium diphtheriae* (PDB ID: 5K9A). The sequence identity between *C. striatum* class C sortase and the selected templates of *Streptococcus pneumoniae, Streptococcus agalactiae, Actinomyces oris, Clostridium perfringens and Corynebacterium diphtheriae* was 50% respectively, which is more than the required 30% sequence homology for generating useful models [40]. As seen in **Figure 1A**, results from I-TASSER modeling of the sortase C protein predicted the presence of all the three secondary structure states (helices, coils, beta-sheets), correlating with the PSIPRED server results **(Suppl Figure 1)**. In Gram-positive bacteria, including *Corynebacterium* spp., the sortase C enzyme is known to possess a highly conserved active site cysteine residue, responsible for cleavage of pilin motifs. Present along with the cysteine residue, are histidine and arginine residues (His and Arg), that play essential roles in sortase activity, thereby forming a catalytic triad [30]. In the I-TASSER model of the *C. striatum* sortase C protein, the triad of Cys230-His168-Arg239 was found between the antiparallel beta-sheet strands **(Figure 1A and B)**. Further, based on molecular surface representation with amino acid hydrophobicity colored based on the Kyte-Doolittle scale [41], the active site is likely in a hydrophobic groove **(Figure 1B)**. The overall quality factor of the predicted sortase C model was 73.0263 by the ERRAT **(Suppl Figure 1)**. Moreover, the model was concurred by VERIFY3D showing 80.45% of the residues with an averaged 3D-1D score ≥ 0.2. Ramachandran plot analysis done by PROCHECK servers revealed that 95.6% of residues of the protein were in the most favored and allowed regions and only 4.4% residues in outlier regions.

**Figure 1:**
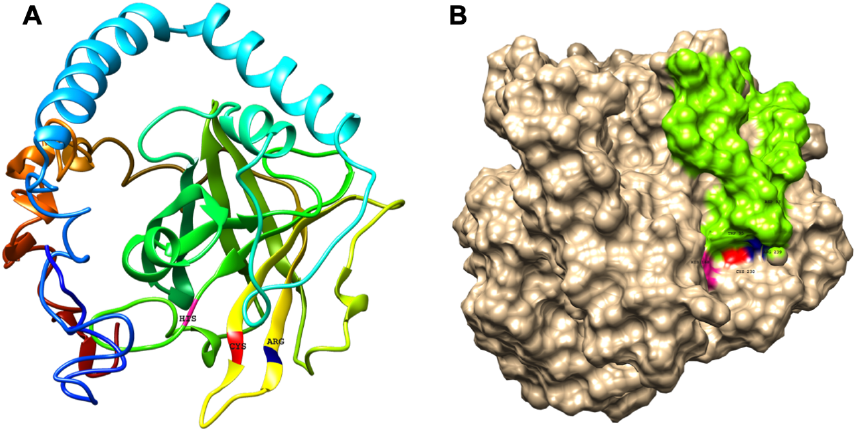
The predicted structure of the *C. striatum* sortase C protein reveals the presence of the putative active site triad, essential for catalytic activity, and a ‘lid’. (A) The sortase C protein in ribbon form, showing the overall organization of the protein backbone in 3D space. The modeled structure reveals the presence of the putative Cys-His-Arg triad essential for catalytic activity. The Cys230 residue is shown in red, the His168 residue in green, and the Arg239 residue in blue. The default ribbon style is smooth (rounded). (B) Surface representation of the class C sortase protein showing the sortase ‘lid’ (green) occluding the active site residues. Homology modeling was done using the I-TASSER server using the amino acid sequence of the sortase C protein retrieved from GenPept (ID: WP_034656562).

### Modeling the *C. striatum* sortase C protein in semi-open lid conformation

In addition to the characteristic beta-barrel structure and active site Cys-His-Arg triad, the homology model of the wild-type sortase C protein predicted the presence of a ‘lid’ consisting of a structural loop lodged in the putative active site. The ‘lid’ of the sortase C protein is suggested to play a role during polymerization of the pilin subunits [33,42], likely via the recognition of the sorting signal. However, there is no evidence that indicates that the pilin substrates are able to access the putative active site in this ‘closed’ confirmation, indicating that the lid likely undergoes displacement during pilus biogenesis [33,42]. As seen in **Figure 2A and B,** the ‘lid’ of the sortase C protein carries two important residues, which typically include an aspartate residue and a hydrophobic amino acid, two residues after aspartate, that interact with the putative catalytic site. In the ‘closed’ confirmation, the sortase C lid blocks access to the active site, observed as proximity between the Asp93 and Arg239 residues. The presence of the aromatic ring of the Trp95 residue in the lid, which is close to the Cys230 residue, renders the active site inaccessible [43]. Point mutations in the ‘lid’, such as a double mutation D93A/W95A, replacing Aspartate (93) and Tryptophan (95) with Alanine, are known to result in a ‘semi-open’ lid conformation and increased substrate accessibility [43]. To mimic the semi-open lid conformation, we generated a model that included a double mutation D93A/W95A in the sortase C protein. As seen in **Figure 2C and D**, I-TASSER modeling of the mutated sortase C protein (D93A/W95A) predicted the presence of a ‘semi-open’ lid conformation, with the lid displaced away from the triad residues and a cavity observed between the alanine residues and the putative active site triad. The mutated sortase C protein displayed similar physicochemical properties and secondary prediction as compared with the wild-type protein **(Table 1)**, with a distribution of alpha-helices (41%), coils (44%), as well as beta sheets (15%) **(Suppl Figure 2)**. Further, the sortase C protein model with ‘semi-open’ lid conformation generated a 3D score of 82.37% by the Verify3D Program under SAVES v6.0. The ‘semi-open’ conformation of the sortase C protein was used for protein-peptide *in silico* molecular docking studies.

**Figure 2:**
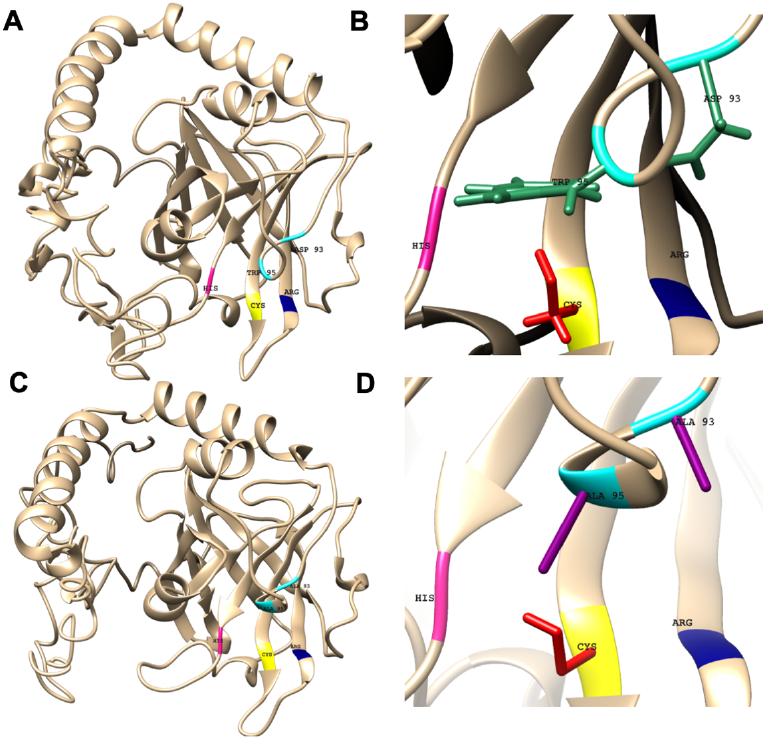
Predicted and chosen models of the wild-type *C. striatum* sortase C protein (closed conformation) and mutant protein (D93A/W95A; semi-open conformation) showing the relationship between the sortase ‘lid’ and putative active site triad residues. (A) In the wild-type sortase C protein, the sortase ‘lid’ is predicted as a loop-like extension, in close proximity with the putative active site Cys-His-Arg (yellow-pink-blue) residues (B) The closer version of the region, predicts interactions between the aromatic ring of the Trp residue (green) and the Asp residue (green) of the ‘lid’ with Cys230 and Arg239 residues respectively (C) In the mutant sortase C protein (D93A/W95A), the lid is predicted to be displaced from the triad residues (D) In the closer version, a cavity is predicted between the mutated alanine residues (purple) and the putative active site triad. This ‘semi-open’ lid conformation of the mutated sortase C protein was used for protein-peptide *in silico* molecular docking studies.

### Predicted structural models of AMPs using sequences from the DRAMP database

A dataset of 5562 general AMP sequences, including natural and synthetic AMPs, were downloaded from the DRAMP V3.0 database [19]. AMPs were listed from Pep2 to Pep5563 (Pep ID), where Pep1 is designated as the pilin subunit (GenPept ID: WP_170219081.1) **(Suppl Fig 3)** [20]. Using a combination of PEP-FOLD3, TrRosetta, and I-TASSER, we developed structural models of 5544 AMPs. Specifically, PEP-FOLD3 was used for peptide sequences with less than 50 amino acids, TrRosetta for sequences with more than 50 amino acids, and I-TASSER for sequences with undetermined ‘X’ amino acids. For 18 AMPs possessing unusual or exotic amino acids, models could not be built as these amino acids were not recognized by the structure prediction tools. Predicted and chosen models were selected based on cluster ranking and scoring. As shown in **Figure 3**, AMPs were predicted to possess a range of structures from alpha helices, beta sheets, coils, or combinations of these. All 5544 predicted and selected peptide structures, searchable by their unique Pep ID, DRAMP ID, name, and course, and a separate subset of AMPs filtered by known anti-Gram positive activity are available in a structural AMP repository for biofilm studies (B-AMP) [44]. For each peptide, the FASTA files, PDB files, and images of the predicted and chosen 3D models are available.

**Figure 3:**
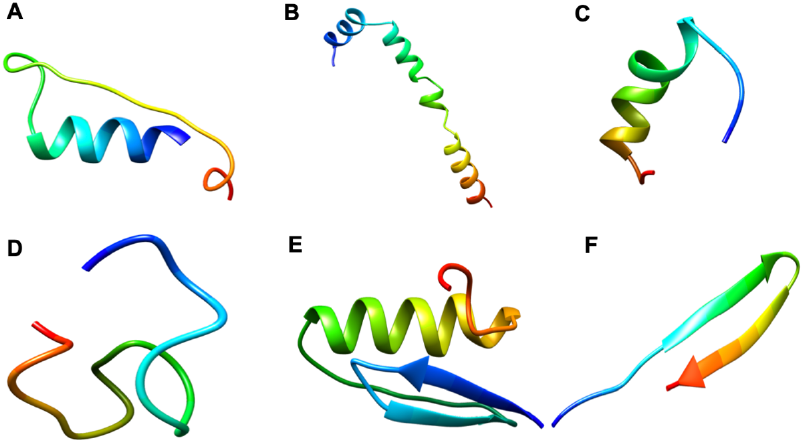
Representative predicted AMP structures with PEPFOLD3, TrRosetta and I-TASSER. FASTA sequences of AMPs from the DRAMP database were modeled using the PEPFOLD3 (for AMPs less than 50 amino acid residues in length), TrRosetta (for AMPs more than 50 amino acid residues in length) and I-TASSER (for AMPs with unusual or unknown amino acids). Predicted AMP structures were observed to possess a range of features such as alpha helices, beta sheets, coils or combinations of these. (A) Pep2 (DRAMP00005) showing a helix-coil structure (B) Pep6 (DRAMP00068) with amino acid sequence showing an alpha-helix structure C) Pep31 (DRAMP00244) showing an alpha-helix structure (D) Pep3 (DRAMP00017) showing a coil structure (E) Pep7 (DRAMP00069) showing an antiparallel beta sheet structure (F) Pep5 (DRAMP00063) showing a combination of alpha helices, beta sheets and coils. The entire structural library of 5544 AMPs is available in B-AMP. Models are colored from blue to red from N-to-C-terminal.

### Protein-peptide docking of candidate AMPs against the mutated sortase C protein of *C. striatum*

To identify candidate AMPs with potential to interact with the putative active site of the sortase C enzyme of *C. striatum*, 88 AMPs with a length ranging from 2-8 residues, and 12 representative AMPs each ranging from 9-20 residues in length were selected. Select peptides were subject to *in silico* molecular docking using AutoDock Vina [29], against the ‘semi-open’ conformation of the mutated sortase C protein. As a comparison, the LPMTG motif of the pilin subunit of *C. striatum* was considered as the standard (where M can be any amino acid). As seen in **Figure 4**, the LPMTG motif interacted with the mutated sortase C protein at the active site, forming three hydrogen bonds with His168, Asn236 and Gln143, with a binding affinity of −5.3 kcal/mol. Of the 100 candidate AMPs, 99 AMPs were predicted to form hydrogen bonds with the mutated sortase C protein (**Figure 4)**. We observed that, for AMPs more than 10 amino acids in length, as the length of the AMP increased the docking score decreased **(Figure 5)**. Further, AMPs more than 14 residues in length did not interact with the putative active site triad residues, but instead masked the active site by binding to protruding residues near the triad. Given this, we decided to analyze specific hydrogen interactions between the 100 candidate AMPs and residues of the mutated sortase C protein **(Figure 5)**. It is important to note that, based on our docking results, no interactions were predicted between the LPMTG motif or any of the AMPs with the active site Cys230 residue. This could be due to the semi-open lid conformation of the sortase C protein that limits access to the Cys230 or could relate to the chemical structure of cysteine that confers it selective interactions. All protein-peptide interaction models are available in B-AMP [44]. For each protein-peptide model, input and output PDBQT files, model images and bond information are provided. It is important to note that AutoDock Vina is typically suited for peptides with less than 32 bonds, and if the number of rotatable bonds exceeds 32 the tool flags select bonds as rigid or non-rotatable to fit parameters. Given this, we decided to work with small AMPs of lengths ranging from 2-8 residues, with a few select AMPs of 9-20 residues in length.

**Figure 4:**
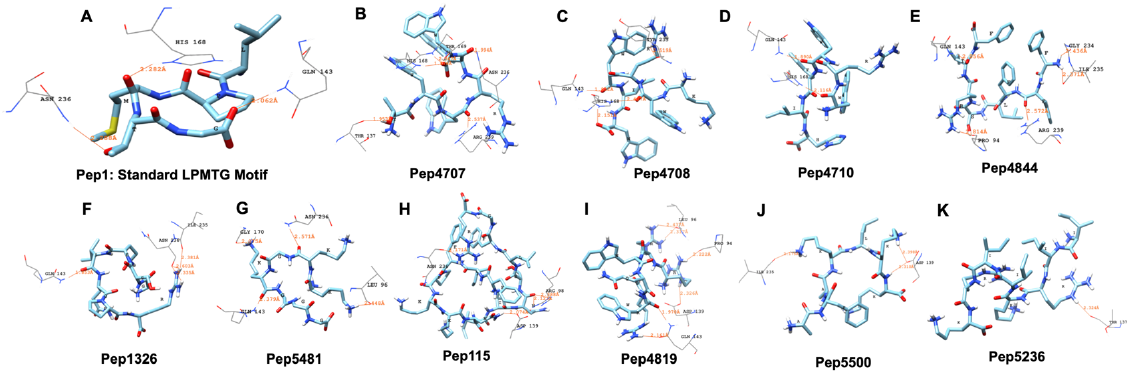
Representative results from *in silico* protein-peptide molecular docking for the standard pilin subunit LPMTG, and AMPs showing a range of hydrogen interactions, with the *C. striatum* mutated sortase C protein. (A) The standard LPMTG motif is predicted to form hydrogen bonds with His168, Asn23y, Gln143 (B) Pep4707 is predicted to interact with HIS168, ARG239, ASN236, THR137, THR169 (C) Pep4708 is predicted to interact with HIS168, HIS168, GLN143, TYR233 (D) Pep4710 is predicted to interact with HIS168, GLN143 (E) Pep4844 is predicted to interact with ARG239, GLN143, PRO94, ILE235, GLY234 (F) Pep1326 is predicted to interact with ASN236, ASN236, GLN143, ILE235 (G) Pep5481 is predicted to interact with ASN236, GLN143, LEU96, GLY170 (H) Pep115 is predicted to interact with ASN236, ASP139, ARG98, ARG98 (I) Pep4819 is predicted to interact with GLN143, GLN143, ASP139, LEU96, LEU96, PRO94 (J) Pep5500 is predicted to interact with ASP139, ASP139, ILE235 (K) Pep5236 is predicted to interact with THR137.

**Figure 5:**
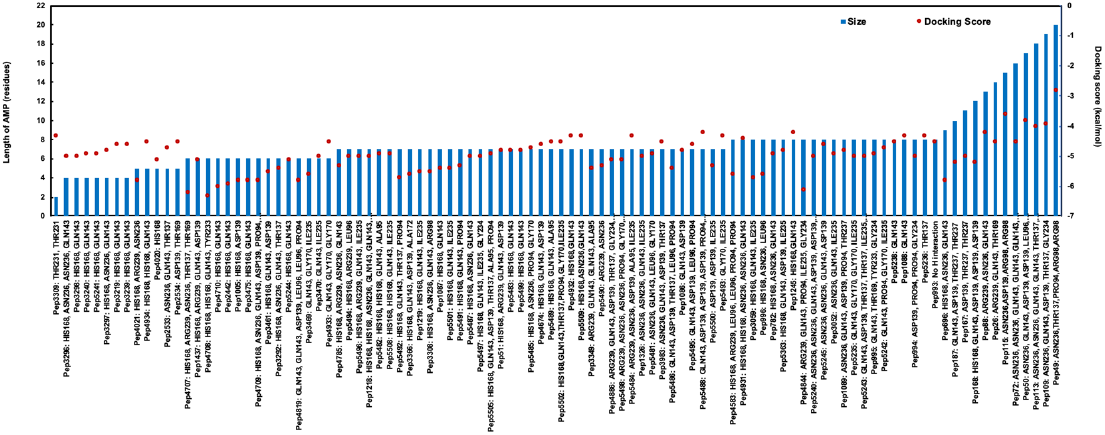
*In silico* molecular docking scores and interacting residues for 100 candidate AMPs and the semi-open (mutated) conformation of the *C. striatum* sortase C protein. AMPs were predicted to interact with the putative active site triad residues, protruding residues near the triad as well as residues further away from the triad. While docking scores were variable, for AMPs more than 10 amino acids in length, as the length of the AMP increased the docking score was observed to decrease.

### Preference scale of candidate AMPs based on docking interactions with mutated sortase C protein of *C. striatum*

Based on *in silico* molecular docking results, we categorized 100 candidate AMPs on a preference scale of 0-10, using interacting residues and docking scores **(Figure 5).** AMPs were categorized, in order of priority starting with interaction with at least two putative catalytic site residues, as seen with His168 and Arg239, which was given a preference score of 10. This was followed by preference score 9, which consisted of AMPs with multiple explicit interactions with His168 from the triad, as well as interactions with other amino acid residues. Similarly, AMPs with only one explicit interaction with His168, along with interactions with other amino acid residues, were given a preference score of 8. AMPs interacting with Arg239 (along with other amino acid residues) were designated a preference score of 7. In the preference score, exclusive interactions with His168 were given priority over Arg239, as the histidine residue is conserved across sortases, whereas arginine can be replaced with another residue. AMPs with interactions with residues surrounding the putative active site triad, such as ASN236 and GLN143, were categorized into preference scores from 6-3. Finally, interactions with ASP139, THR137, GLY170, ILE235, THR231 were categorized into preference scores from 2 and 1, based on their location further away from the triad. A single AMP showed no hydrogen bond interactions with the sortase C protein, and was given a preference score of 0. Finally, in each preference score, candidate AMPs were ordered based on docking scores, from highest to lowest scores **(Table 2)**.

**Table 2:**
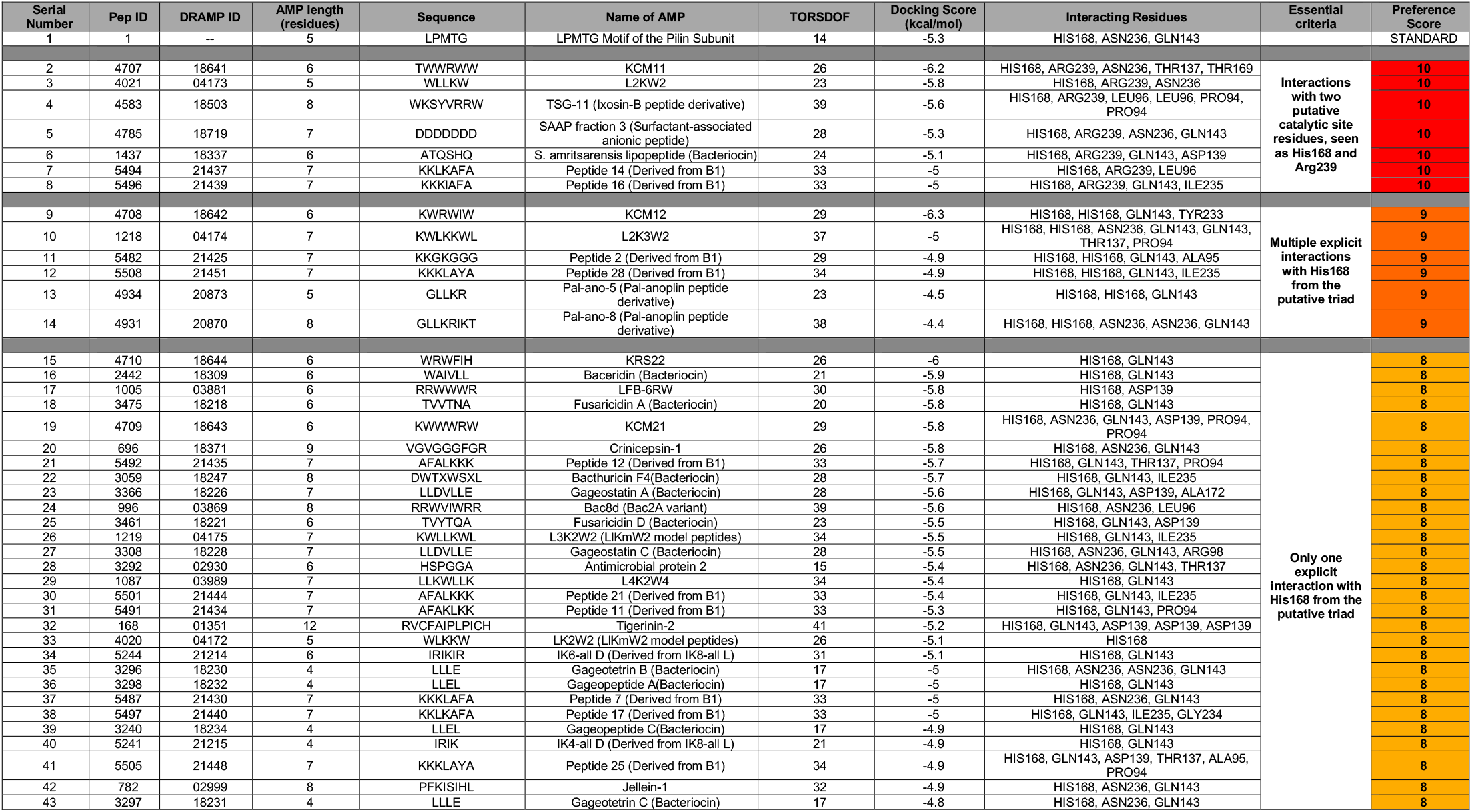
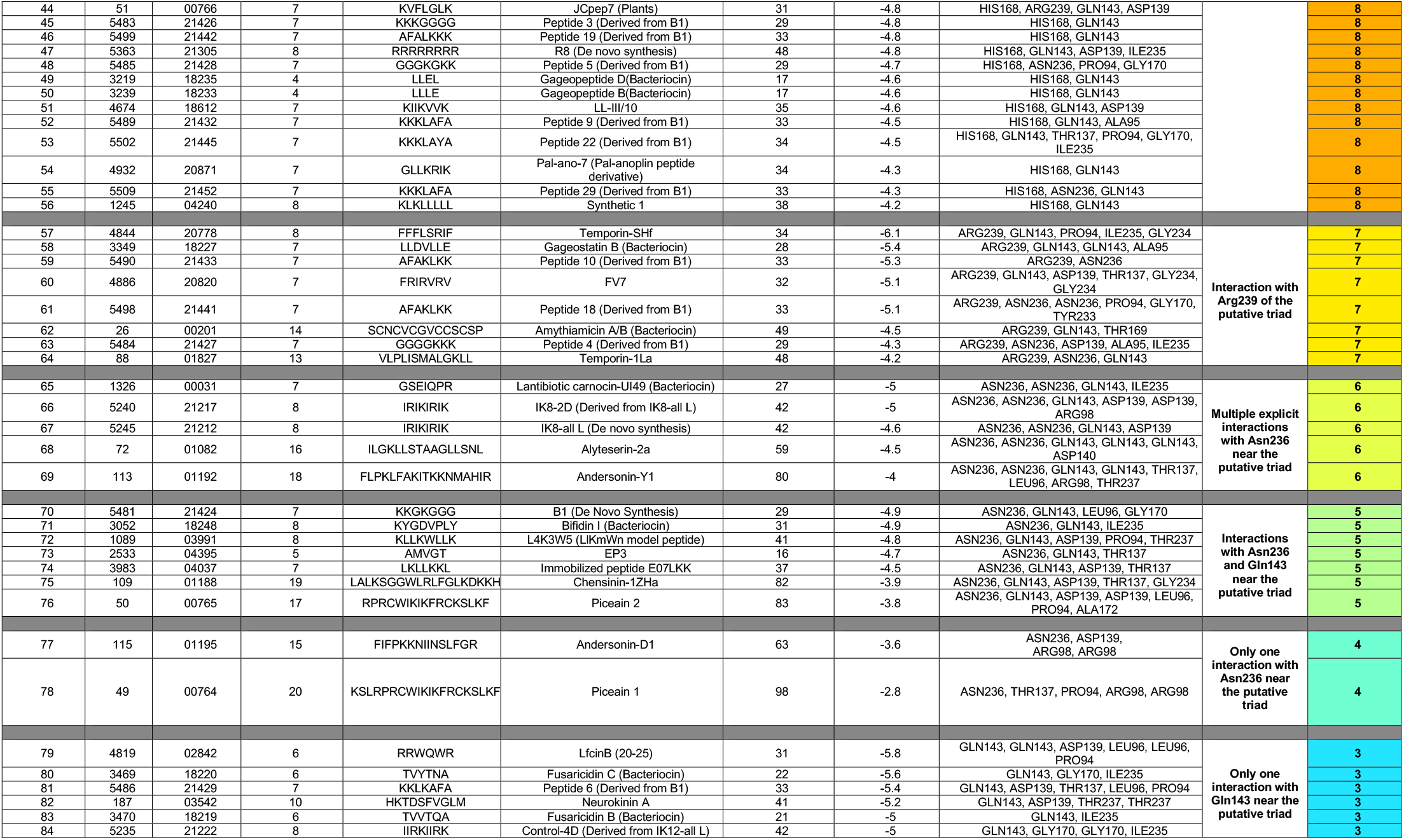
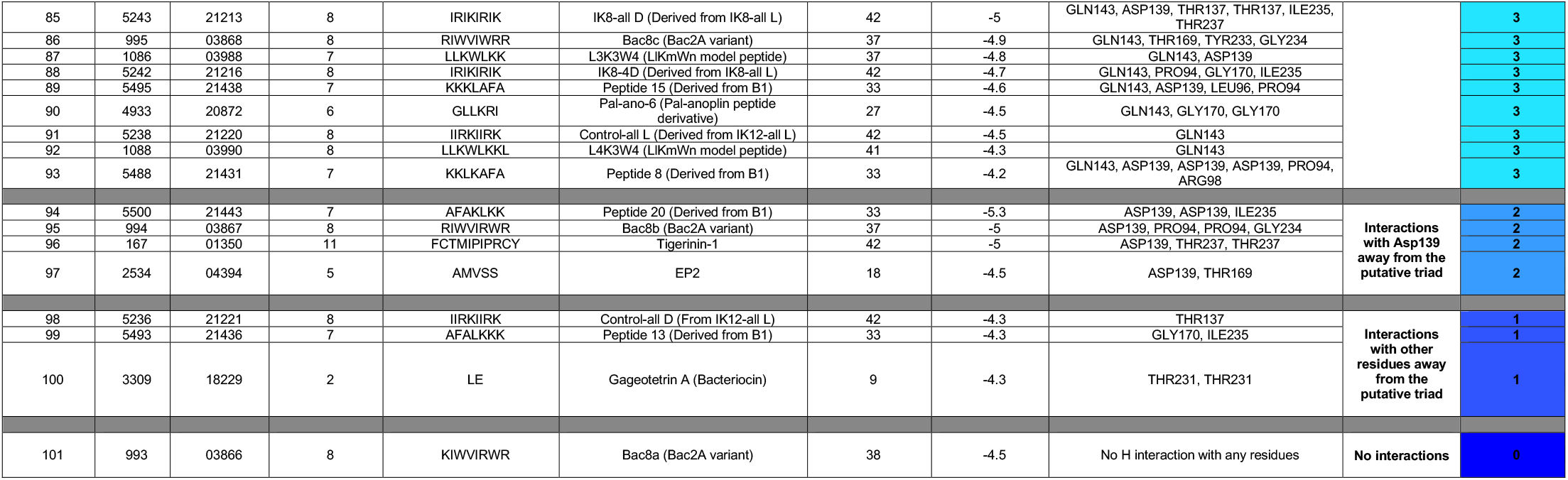
Preference scale of candidate AMPs based on docking interactions with mutated sortase C protein of *C. striatum*. (where 10 represents the most optimum docking interactions and 0 represents no interactions with the protein residues)

### Biofilm-AMP (B-AMP)

The resources and data generated in this approach are freely available in Biofilm-AMP (B-AMP), a user-friendly, search-enabled repository of AMP structures and AMP interactions with biofilm targets (https://b-amp.karishmakaushiklab.com/). The repository includes predicted structures for 5544 AMPs, searchable by their Pep ID, DRAMP ID, name and source, and a separate subset of 2534 AMPs filtered by known anti-Gram positive activity. For each peptide, the repository includes FASTA files, PDB files, and predicted and chosen 3D models. For the 100 AMPs selected for *in silico* molecular docking, the repository hosts input & output PDBQT files, images, and a text file detailing hydrogen interactions **(Figure 6)**. The Python scripts used for auto-generating FASTA files and filtering out peptides are available in a GitHub repository (https://github.com/KarishmaKaushikLab/B-AMP). B-AMP will be continually updated with new structural AMP models and AMP interaction models with potential biofilm targets.

**Figure 6:**
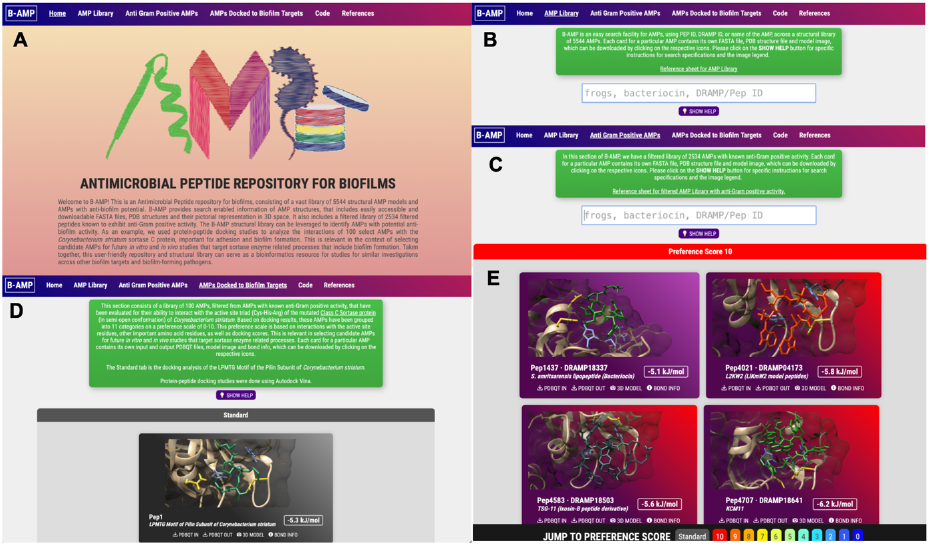
Features of Biofilm-AMP (B-AMP). (A) B-AMP is a structural repository of AMPs for biofilm studies (B) This user-friendly, search enabled repository includes predicted structures for 5544 AMPs, searchable by their Pep ID, DRAMP ID, name and source (C) It also includes a separate subset of 2534 AMPs filtered by known anti-Gram positive activity. For each peptide, the repository includes FASTA files, PDB files, and predicted and chosen 3D models (D and E) For the 100 AMPs selected for *in silico* molecular docking, the repository hosts protein-peptide docking models, input & output PDBQT files, images, and AMPs categorized in order of preference, based in docking scores and interacting residues. B-AMP will be continually updated with new structural AMP models and AMP interaction models with potential biofilm targets.

## Conclusions

AMPs are increasingly being recognized for their anti-biofilm potential, to act as alternatives to or in adjunct with conventional antibiotics [45–48]. Their ability to target specific and conserved cellular processes, hold particular promise in the context of broadspectrum activity and decreased possibility of resistance development [47]. While several AMPs have been studied for their ability to inhibit biofilm formation [49], it is imperative to enable a steady pipeline of candidate AMPs for anti-biofilm evaluation. However, given the vast array of natural and synthetic AMPs, identifying potential candidates for *in vitro* and *in vivo* anti-biofilm testing remains a significant challenge.

Our approach uses open-source bioinformatics tools, to identify AMPs with potential anti-biofilm activity, using the sortase-pilin machinery of *C. striatum* as the target **(Figure 7).** With *in silico* protein-peptide molecular docking, we identified AMPs predicted to interact with putative catalytic site residues of the *C. striatum* sortase C protein, or residues in proximity with the triad. This could result in reduced accessibility of the pilin substrate to the catalytic site or inhibition of catalytic function of the protein [31,50,51]. Based on our approach, we propose a preference scale, based on which candidate AMPs can be taken up, in order of priority, for subsequent evaluation of anti-biofilm activity against *C. striatum*. This can include additional *in silico* approaches such as molecular dynamics simulations and RMSD analyses, in addition to standard *in vitro* and *in vivo* anti-biofilm testing approaches, such as assays for metabolic activity, biofilm eradication and inhibitory concentrations, advanced microscopy, and animal models of infection [51]. Further, evaluations can also focus on bioactivity, stability, and toxicity testing of candidate AMPs. To improve biocompatibility and minimize toxicity, candidate AMPs can also be subject to peptide design and engineering to produce suitable modifications [47,52–54]. While certainly not inclusive of all aspects critical to the evaluation of candidate AMPs as potential anti-biofilm agents, our approach uses open-source bioinformatics tools to narrow down candidate AMPs from a vast library of natural and synthetic AMPs. It thereby provides a starting point for subsequent *in silico*, as well as *in vitro* and *in vivo* biofilm testing approaches against an emerging, multidrug resistant, biofilm-forming pathogen.

**Figure 7:**
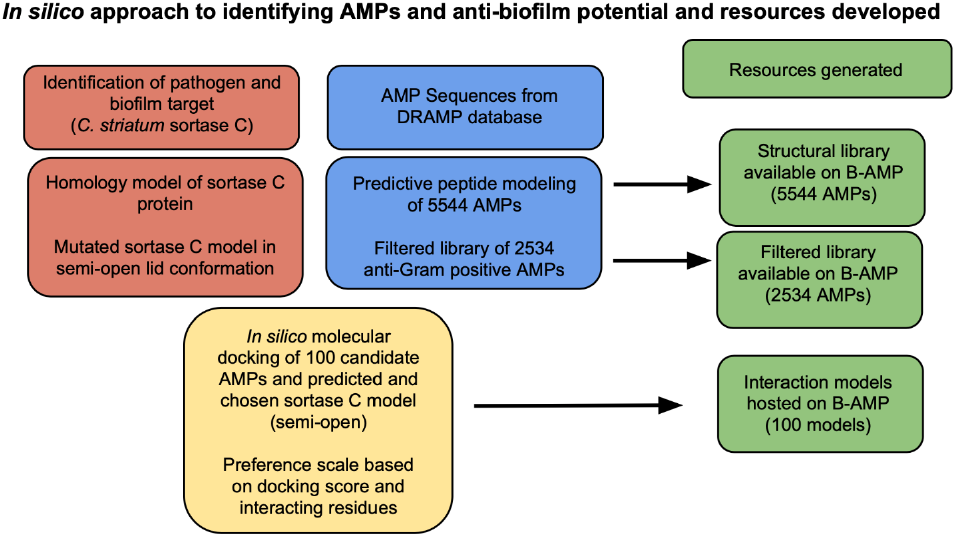
*In silico* approach to identifying AMPs with anti-biofilm potential and resources developed in this study. Our approach used a combination of homology modeling, predictive peptide modeling and protein-peptide molecular docking. The resources and data generated in this approach are freely available in Biofilm-AMP (B-AMP), a user-friendly, search-enabled repository of AMP structures and AMP interactions with biofilm targets. The approach used and resources developed can be leveraged for similar studies against other biofilm targets and biofilm-forming pathogens.

While our study focused on the sortase-pilin machinery of *C. striatum*, the approach used and resources developed, which includes a unique structural repository of AMPs and protein-peptide models, can be leveraged for similar studies against other biofilm targets and biofilm-forming pathogens. Further, the vast structural library of AMPs in B-AMP, can be leveraged for a range of AMP-related investigations across a range of bacterial and fungal species.

## Supplementary figures

**Suppl Fig 1:**
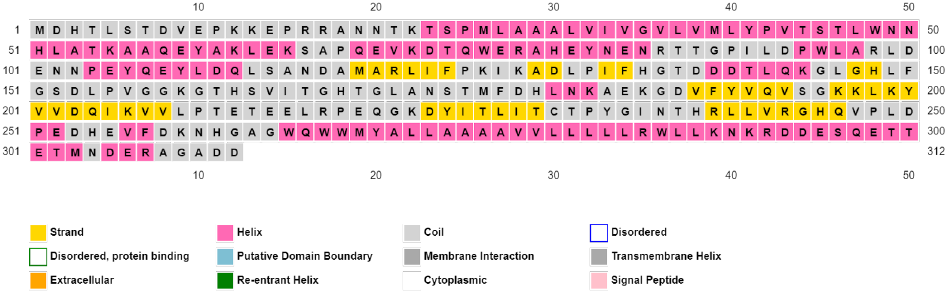
Secondary structure of the sortase C protein based on PSIPRED analysis. The secondary structure of the sortase C protein contains a combination of alpha-helices (41%), coils (44%), and beta strands (15%). The cysteine (Cys230) and histidine (His168) residues are found to be in between two beta-strands, constituting a turn connecting the strands and acting as a major groove for hydrophobic interaction. The arginine (Arg239) residue was predicted to initiate the second beta-strand of the hydrophobic groove. The image is a snapshot of the results obtained from the PSIPRED analysis program.

**Suppl Fig 2:**
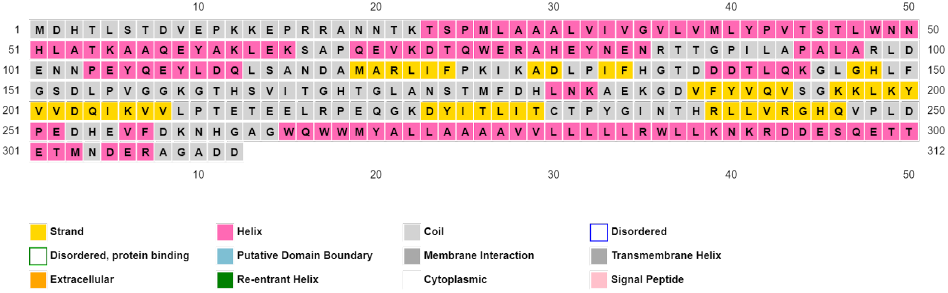
Secondary structure of the mutated sortase C protein based on PSIPRED analysis. The secondary structure of the mutated sortase C protein with Aspartate93 and Tryptophan95 replaced with Alanine (D93A/W95A). The image is a snapshot of the results obtained from the PSIPRED analysis program.

**Suppl Fig 3:**
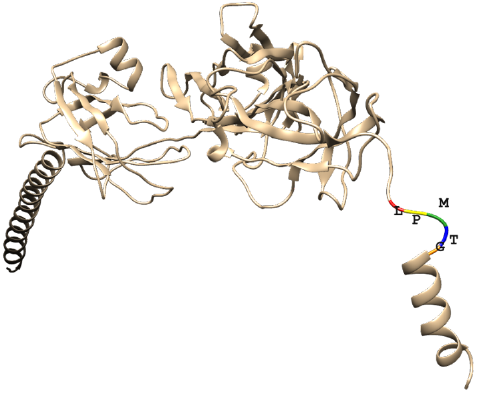
Predicted and chosen model of the *C. striatum* pilin subunit showing the presence of the cell sorting LPMTG motif. The *C. striatum* pilin subunit in ribbon form, showing the overall organization of the protein backbone in 3D space. The modeled structure rivals the presence of the cell sorting LPXTG motif (X is represented by M) at the C terminus of the protein. This motif is followed by a hydrophobic domain seen here as a helix. Homology modeling was done using the I-TASSER Server using the amino acid sequence of the pilin subunit retrieved from GenPept (ID: WP_170219081.1).

## Funding

Karishma S Kaushik’s academic appointment is funded by the Ramalingaswami Re-entry Fellowship (BT/HRD/35/02/2006). Snehal Kadam was supported for a select duration of this project (June 2020-December 2020) on the Ramalingaswami Re-entry Fellowship (to KSK).

## Acknowledgments

We thank Dr. Ragothaman Yenamalli, SASTRA University, Thanjavur, India, for inputs related to *in silico* molecular docking.

## Author Contributions

**Shreeya Mhade:** Conceptualization, Methodology, Investigation, Validation, Formal analysis, Data curation, Visualization, Writing the original draft, Editing draft. **Stutee Panse:** Conceptualization, Methodology, Investigation, Validation, Formal analysis, Data curation, Visualization, Writing the original draft, Editing draft. **Gandhar Tendulkar:** Conceptualization, Methodology, Investigation, Validation, Formal analysis, Data curation, Visualization, Writing the original draft, Editing draft. **Rohit Awate:** Methodology, Investigation. **Snehal Kadam:** Conceptualization, Investigation, Editing draft. **Karishma S Kaushik:** Conceptualization, Project administration, Supervision, Writing the original draft, Editing draft.

## Conflict of Interest

None

## Notes

### Competing Interest Statement

The authors have declared no competing interest.

### Summary of Updates

Figure 5 - was accidentally cut off during upload.

